# Using inbreeding to test the contribution of non-additive genetic effects to additive genetic variance: a case study in *Drosophila serrata*

**DOI:** 10.1101/2023.01.22.525104

**Authors:** Robert J. Dugand, Mark W. Blows, Katrina McGuigan

## Abstract

Additive genetic variance, *V_A_*, is the key parameter for predicting adaptive and neutral phenotypic evolution. Changes in demography (*e.g*., increased close-relative inbreeding) can alter *V_A_*, but how depends on the, typically unknown, gene action and allele frequencies across many loci. For example, *V_A_* increases proportionally with the inbreeding coefficient when allelic effects are additive, but larger (or smaller) increases can occur when allele frequencies are unequal at causal loci with dominance effects. Here, we describe an experimental approach to assess the potential for rare, recessive alleles to inflate *V_A_* under inbreeding. Applying a powerful paired pedigree design in *Drosophila serrata*, we measured 11 wing traits on half-sibling families bred via either random or sibling mating, differing only in homozygosity (not allele frequency). Despite close inbreeding and substantial power to detect small *V_A_*, we detected no deviation from the expected additive effect of inbreeding on genetic (co)variances. Our results suggest the average dominance coefficient is very small relative to the additive effect, or that allele frequencies are relatively equal at loci affecting wing traits. We outline the further opportunities for this paired pedigree approach to reveal the characteristics of *V_A_*, providing insight into historical selection and future evolutionary potential.

## Introduction

Adaptive evolution, and the maintenance of fitness in natural populations, depends on the distribution of phenotypes, and on the presence and nature of genetic variation underlying those phenotypes [1]. The most important metric of genetic variation for complex phenotypic traits is additive genetic variation (*V_A_*), which causes most of the resemblance amongst relatives, and, thus, the opportunity for evolution [2–4]. *V_A_* is determined by the number of loci affecting a trait, their allele frequencies, and the effects of those alleles, which can collectively be considered the genetic architecture. Hence, *V_A_* observed for any trait has been determined by evolutionary forces that alter the frequencies of alleles, and, consequently, the frequency distribution of allelic effects. As a summary statistic, the magnitude of *V_A_* does not provide insight into the underlying genetic architecture, but this architecture can affect rates of both phenotypic evolution and depletion of evolutionary potential [5].

Importantly, it is not only the additive effects of alleles that determine *V_A_*, but also the non-additive, interaction, effects within and among loci [2, 6, 7]. Indeed, the relative magnitudes of genetic variance components (additive, dominance, or epistatic) provide no insight into the underlying gene action [6]. The contribution of dominance effects to *V_A_* may be particularly interesting as habitat loss and increasingly widespread physical barriers to dispersal increase the frequency of inbreeding (close-relative mating) within populations. In the presence of exclusively additive gene action, inbreeding results in the redistribution of variance among and within families, as well as increasing the total genetic variance at a rate of 1 + *F*, where *F* is the inbreeding coefficient [8, pp. 264-265]. However, when there is dominance, any change in *V_A_* with inbreeding depends on the complex relationship between gene action and allele frequency (**Figure 1**). For example, inbreeding will inflate *V_A_* above (1 + *F*)*V_A_* when alleles are rare (*e.g., q* < 0.05) and recessive (*d* → *a*), but *V_A_* will be less than (1 + *F*)*V_A_* with inbreeding if recessive alleles are common (*e.g*., *q* > 0.95; **Figure 1**).

**Figure 1.**
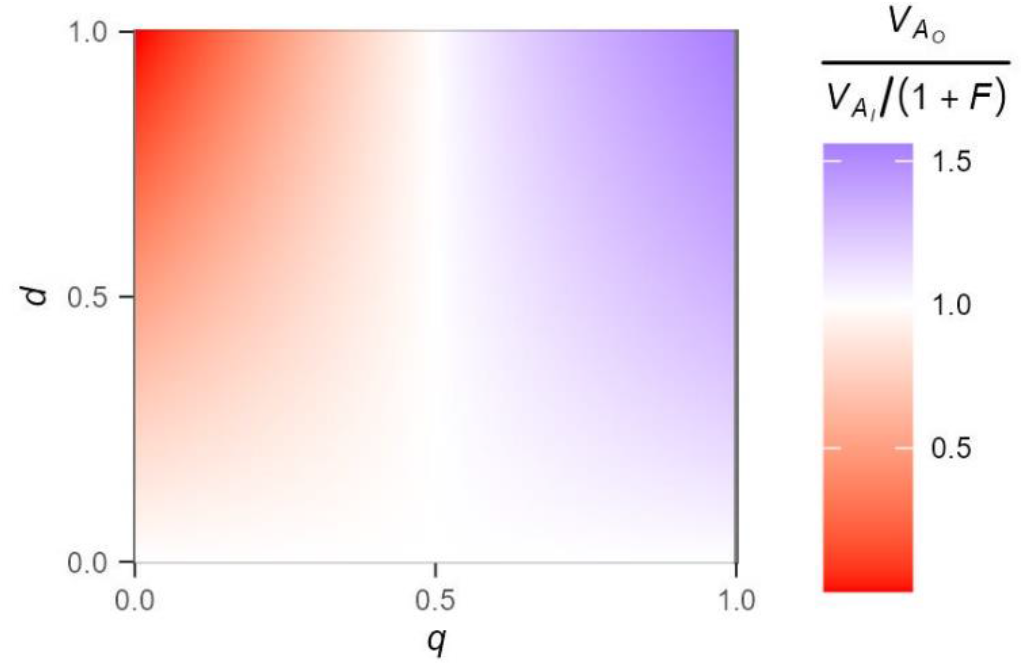
Heat map showing the relative magnitude of additive genetic variance (*V_A_*) in the absence (*V_A_O__*) *versus* presence (*V_A_I__*) of inbreeding. For a given locus with two alleles, the magnitude of *V_A_* can be defined as [19]:

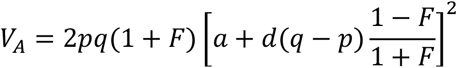

where *p* and *q* are allele frequencies, *F* is the inbreeding coefficient, *a* is the additive effect, and *d* is the dominance coefficient. *V_A_* was estimated from this equation, assuming *a* = 1, and *F* = 0 or *F* = 0.25 (*e.g*., brother-sister inbreeding), and varying *d* from 0 (no dominance) to 1 (*d* = *a; i.e*., complete dominance), and the frequency of the recessive allele (*q*) from 0 to 1. The effect of *d* and *q* are illustrated by plotting the relative magnitude of *V_A_* in the absence or presence of inbreeding, calculated as 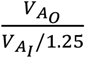, thus accounting for the increase in *V_A_* that is expected under a purely additive genetic model. Red = *V_A_I__*/(1 + *F*) > *V_A_O__*; white = *V_A_I__*/(1 + *F*) = *V_A_O__*; blue = *V_A_I__*/(1 + *F*) < *V_A_O__*.

Approaches such as Genome-Wide Association Studies (GWAS) have revealed that quantitative trait variation is typically strongly influenced by common (*q* > 0.1) alleles across multiple loci, but GWAS are limited in their power to identify rare variants, or those with complex (non-additive) effects [9]. Large-scale studies, particularly in humans, are starting to overcome these limitations. Critically, rare alleles have been shown to contribute substantially to variation of diverse traits, including gene expression and size [10, 11]. Allele frequency has been shown to be negatively correlated with effect size [12–14], suggesting that alleles with larger effects on the focal trait are more deleterious and kept at lower frequencies by (purifying) selection [15]. Such alleles are also hypothesised to be recessive or partially recessive – deleterious alleles with additive and dominant effects will be more rapidly removed by selection [5]. Furthermore, loci at which alleles affecting a phenotypic trait have a pleiotropic effect on fitness (as suggested by the bias toward lower frequency of larger effect alleles) are predicted to contribute to increased variance under inbreeding due to dominance effects [16]. Therefore, although the distribution of dominance coefficients across loci remains poorly resolved [17, 18], prevailing evidence implies a major contribution of rare, (partially) recessive alleles to variation across a range of quantitative traits, and suggests that *V_A_* will be inflated above (1 + *F*)*V_A_* with inbreeding.

The question of how the genetic architecture may influence the magnitude of *V_A_* under elevated inbreeding must be considered from a multivariate perspective. Although most individual quantitative traits have abundant *V_A_* [20], when multiple traits are considered together, one multi-trait combination typically accounts for much of the total *V_A_*, leaving other multi-trait combinations with low *V_A_* [21]. There is evidence that multi-trait combination with low *V_A_* are correlated with fitness [22–24], and determined by rare alleles [25]. If generally true, inbreeding may disproportionately inflate *V_A_* in such multi-trait combinations, which, in turn, could be critical for the maintenance of fitness in current conditions or adaptation to changing environmental conditions.

Here, we applied inbreeding treatments in an outbred population of *Drosophila serrata* to test how changing genotype, but not allele, frequencies affected the distribution of additive genetic (co)variances (**G**) for size and a set of 10 wing shape traits. *Drosophila* wings have emerged as a powerful model in evolutionary genetics [26, 27]. In common with other trait types, the magnitude of *V_A_* varies markedly across different multivariate wing traits in *D. melanogaster* [28] and *D. serrata* [29, 30]. Sztepanacz and Blows [30] detected statistically significant additive and dominance (*V_D_*) genetic variance for several of these wing traits in *D. serrata*. Intriguingly, they observed that the ratio, *V_A_: V_D_*, varied across multi-trait space and was relatively high for the multivariate wing trait under the strongest selection [30]. Because non-additive gene action contributes to *V_A_* when allele frequencies are uneven [7], this observation is consistent with dominance effects at (some of the) loci at which selection has caused allele frequencies to be uneven. Here, we interrogate the effects of inbreeding on the distribution of *V_A_* to gain insight into the genetic architecture of wings (see **Figure 1**), and characterise how evolutionary potential may be affected under conditions of increased close-relative inbreeding.

Our experimental outbred population of *D. serrata* was maintained using a breeding design that maximises the effective population size and minimises genetic drift and inbreeding [31]. Our goal was to understand whether dominant gene action contributed to additive genetic (co)variance, and how such contributions were distributed across multivariate trait space and, in particular, whether variable genetic architecture would result in inbreeding disproportionately inflating *V_A_* for multi-trait combinations that have little evolutionary potential (low *V_A_*) in the outbred population. We estimate **G** in the presence *versus* absence of brother-sister mating, based on a shared population of sires, using the animal model framework [32]. This model accounts for the increase in total variance (1 + *F*) predicted under inbreeding when effects are purely additive [19]. Any divergence in **G** between inbred and outbred populations is, therefore, interpreted as the contribution of dominant gene action, causing the inbred **G** to depart from the predicted 1 + *F* change in variance (**Figure 1**).

## Materials and Methods

### Experimental overview

The experimental design and data collection protocols follow those described elsewhere [29] for the outbred population (defined below). Briefly, from a laboratory-adapted (80 generations of random mating in bottle culture at *N*~3,000) population of *D. serrata*, we established a middle-class neighbourhood [MCN, 33] population consisting of 600 males and 600 females per generation, maintained for 14 generations. Each generation, 600 pairs (haphazardly paired, avoiding any brother-sister pairs) were mated and a single son and daughter from each pair was then used to propagate the next generation, again by random, outbred, pairings. Sires from these crosses, the focal males, were then paired with their sister to produce inbred offspring (**Figure 2A**). Each generation, we collected offspring from each of the ^~^1,200 crosses, resulting in phenotypes of flies that were the product of matings of either unrelated individuals (‘outbred’ offspring henceforth; *F*~0) or full siblings (‘inbred’ offspring henceforth; *F*~0.25). The MCN design maximises the effective population size and minimises genetic drift, thus reducing the probability that rare alleles will be lost through sampling effects.

**Figure 2.**
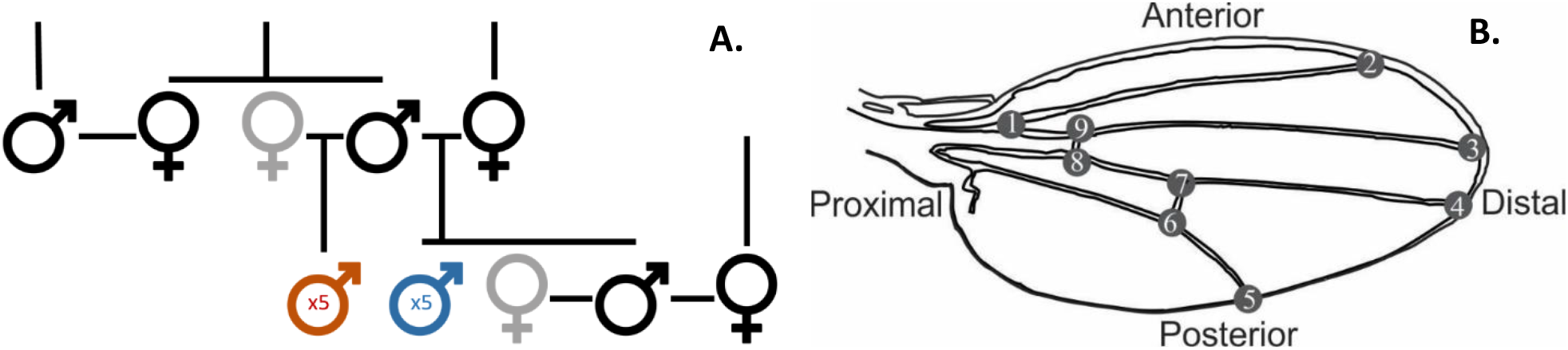
Overview of the experimental design. **A**. Schematic of the pedigree design, showing one family unit across two generations. Focal individuals, which contributed genes to the next generation, are shown in black; females contributing only non-continuing, inbred, offspring are shown in grey. Wing data were collected for up to five inbred (red) and five outbred (blue) males per focal sire each generation. **B**. Schematic of a *Drosophila* wing, depicting the positions of the nine recorded landmarks. Wing size was characterised as centroid size, CS, the square-root of the sum of squared distances between the centroid and each aligned landmark [34]. Following alignment, and accounting for variation in CS, distances were calculated between pairs of landmarks in units of CS. These inter-landmark distance (ILD) traits are referred to by the defining landmarks; *e.g*., ILD1.5 is the distance between the first and fifth landmarks.

By using the same sires, and dams drawn from the same population, at the same frequency (one per family) in the outbred and inbred crosses, this design constitutes parallel populations with equivalent allele frequencies, but different genotype frequencies. Assuming Hardy-Weinberg Equilibrium, when the frequency of the rare allele (*q*) at a locus is below 1%, these rare alleles will be present in homozygous individuals at least 25 times more frequently in the presence of inbreeding (**Figure 3**), substantially increasing the potential for rare recessive alleles to contribute to trait variation. As the frequency of alleles at a locus becomes more equal (*i.e., q* → 0.5), the relative frequency of homozygous individuals is increasingly comparable in the presence and absence of inbreeding (**Figure 3**). Comparison of **G**, which captures the cumulative change in genotype frequency across all contributing loci, therefore, provides a test of the contribution of rare, recessive alleles to trait variation.

**Figure 3.**
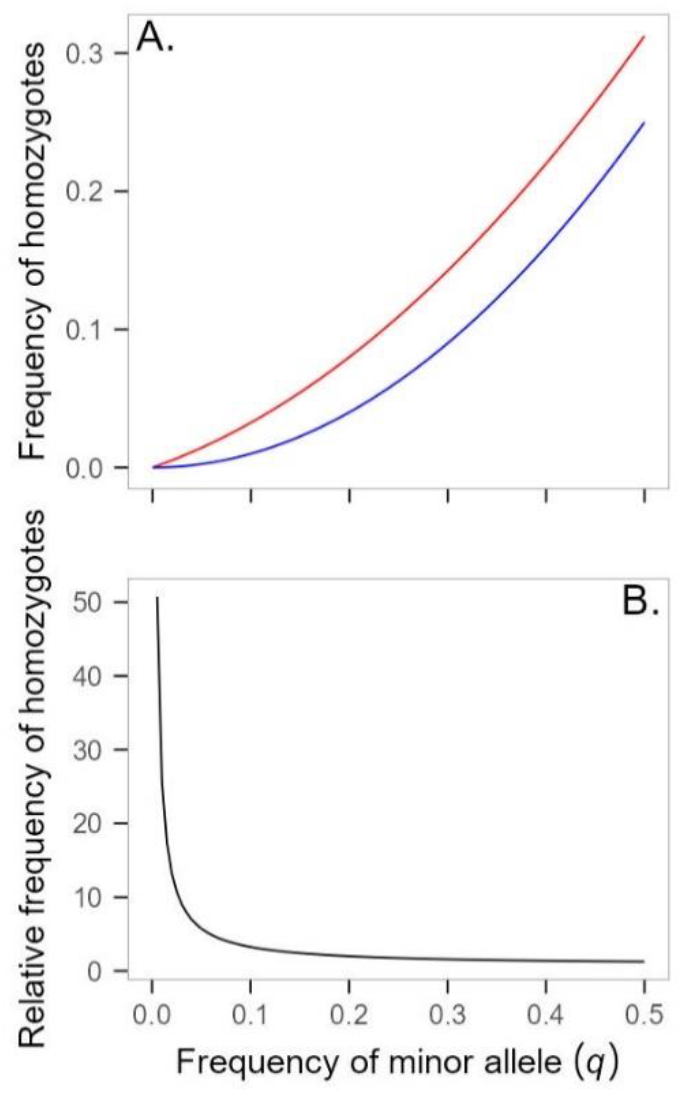
The effect of close inbreeding on the frequency of homozygotes in a population and consequently on the potential for dominant gene action to contribute to trait variation. **A**. The frequency of individuals that are homozygous for allele, *q*, in the presence (*F* = 0.25; *q*^2^ + 0.25[*q*(1 – *q*)]; red line) or absence of inbreeding (*F* = 0; *q*^2^; blue line), plotted as a function of the allele frequency, assuming Hardy Weinberg equilibrium. **B**. The frequency of individuals homozygous for allele, q, in the presence of brother-sister inbreeding relative to their frequency in the absence of inbreeding, plotted as a function of allele frequency.

We collected wing data from up to five non-focal males per cross per sire (**Figure 2**). Wings (one wing per male) were removed, mounted on slides, and photographed in groups of 10-20. Due to changes in data collection protocols, wings from generations 1-4 were unsuitable for inclusion, and here we analyse wing phenotypes from generations 5 through 14; the pedigree information for all generations was included. Photos were randomised and the wings landmarked by one of six observers. Co-occurrence within a single image meant that the wings for a given full-sib family were photographed and landmarked together, which could inflate the resemblance among brothers. However, other relationships, within and across generations, would be unaffected, and any inflated resemblance among brothers is captured by the among-vial variance (see **equation 1**).

We used *tps* software [35] to locate nine landmarks on the image of each wing (**Figure 2B**). Landmarks were then aligned using full Procrustes fit in MorphoJ [36]. Centroid size and aligned XY coordinates were recorded. We then calculated inter-landmark distances (ILDs) for 10 traits which, together with centroid size (wing size henceforth), were the focus of the following analyses (see **Figure 2** for trait descriptions).

After transforming each observation to a deviation from the mean for their respective level of trait, observer, and generation, we identified and removed 449 (1.06% of the data) outlier wings based on Mahalanobis distance (*χ^2^* critical value for α = 0.001 and d.f. = 11). We then filtered the data to retain only phenotypes where sires contributed both inbred and outbred offspring. That is, the phenotype of a given male was only included in the analysis if he had at least one half-brother that had also been assayed. A total of 31,457 wings (15,335 outbred, 16,122 inbred) from 3,502 sires were included in the analysis (the pedigrees included relationship information from a further ^~^4,200 sires whose phenotypes were excluded for the reasons detailed above). Trait values were multiplied by a constant to facilitate model convergence (ILD traits were x1,000, wing size was x200). While data were mean-centred to identify outlier wings, the analyses (detailed below) were conducted on the raw, not mean-centred data.

### Data analysis and parameter estimation

Rare, recessive alleles segregating in a population are expected to consistently reduce fitness, such that inbreeding causes a directional change in trait mean; this pattern is typically referred to as inbreeding depression (ID), and ID is commonly used to diagnose the presence of loci with dominant gene action. However, while the presence of ID directly demonstrates dominant gene action, the absence of ID does not preclude dominant gene action. For traits under stabilising selection, approximately half of the rare, recessive, deleterious alleles will increase trait values, while the other half will decrease trait values [37]. Thus, non-additive gene action can contribute to *V_A_* and changes in *V_A_* under inbreeding, but not result in directional shifts in trait mean of inbred individuals.

We calculated the inbred and outbred offspring means for each of the 3,502 sires and calculated ID for each sire family as: 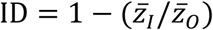, where 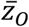 is the outbred mean and 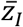 is the inbred mean assuming offspring are fully inbred (*F* = 1), calculated via linear extrapolation [Table 10.2, 38]. Thus, ID is the proportional change in trait value due to inbreeding.

To estimate additive genetic (co)variances, we performed two multivariate analyses, one including only the outbred phenotype data, and one including only the inbred phenotype data. Wombat [39] was used to fit the model:

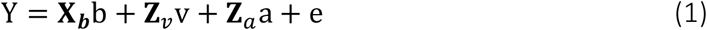

where Y is a vector of phenotypic records, **X_*b*_** is the design matrix relating phenotypic records to the vector of fixed effects, b, that included trait, generation, and observer, and *e* is the residual error. **Z***_v_* and **Z***_a_* are design matrices that relate records to the random vial and additive genetic effects. The vectors v, a, and e denote the predicted vial, additive genetic and residual deviations for each individual for each trait and have associated covariance matrices equal to 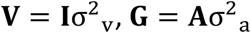, and 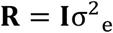, where **A** is the numerator relationship matrix and **I** is the identity matrix. We use the subscripts “O” and “I” to designate the parameters estimated in the analysis of the Outbred and Inbred data, respectively, *i.e*., **G_O_** and **G_I_.**

In the presence of exclusively additive gene action, inbreeding increases the genetic variance at a rate of 1 + *F* [8, pp. 264-265]. The increase in *V_A_* is readily accounted for by **A**, where, for example, the diagonals of **A** equal 1 + *F* [32]. This statistical correction for inbreeding means that, in the absence of non-additive gene action, **G_O_** and **G_I_** will be equal (given sampling error; **Figure 1**, where *d* = 0). To assess the hypothesis that non-additive gene action alters the effect of inbreeding on additive genetic (co)variance, our hypothesis tests focus on **G_O_** = **G_I_**.

#### REML-MVN sampling

We used REML-MVN sampling on the G-scale [40, 41] to place confidence intervals around the estimates and metrics derived from them (*e.g*., eigenvalues). The REML estimates of **G_O_** and **G_I_** were constrained to be positive-definite (*i.e*., no negative estimates of trait variance and no negative eigenvalues), but the REML-MVN samples of this parameter space are unconstrained under the G-scale sampling approach [41]. The parameter estimates and average information (AI) matrix are output from Wombat. We inverted the AI matrix in R [42] using the *solve* function, and used the *mvrnorm* function in the MASS package [43] to estimate 10,000 samples for each **G** from the multivariate normal distribution (*mu*, *Sigma*), where *mu* is a vector of parameter estimates and *Sigma* is the inverse of the AI matrix. We conducted all comparisons between the two **G** based on estimates from the full-rank models, allowing us to probe similarity across the entire multivariate trait space. We used R [42] for all matrix comparisons, with figures produced using *ggplot2* [44].

#### *Comparing* G

First, we investigated how inbreeding affected the size (magnitude) of *V_A_* (**Figure 4A**). Genetic variances (the diagonals of **G**) provide insight into the evolutionary potential of individual traits, with the trace (the sum of the variances) summarising the total genetic variance (*i.e*., matrix size), and evolutionary potential of the total trait set. To statistically test for differences in variances, we paired REML-MVN samples and calculated the ratio of variances (*V_A_O__/V_A_I__*) for each trait in each pair of samples, as well as the total genetic variance. We inferred statistical support for a departure from the (1 + *F*)*V_A_* prediction when 95% CI of the ratio did not overlap 1.0.

**Figure 4.**
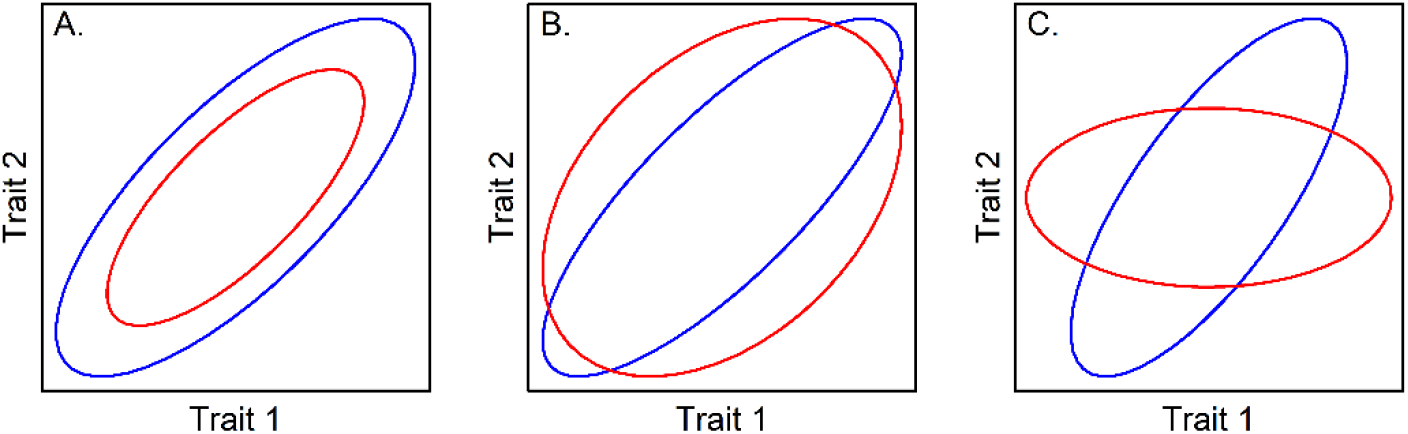
Schematic representation of the approaches for comparing **G**, and their interpretation. Two G-matrices (red and blue) are plotted as ellipses, representing the distribution of genetic variation among two traits. We compared **G** in terms of their size (**A**), shape (**B**), and orientation (**C**).

We next evaluated whether the shape of **G** was affected by inbreeding (**Figure 4B**), taking several approaches. First, we compared the genetic covariance among trait pairs and determined statistical support for differences when the 95% CI did not overlap. Genetic covariance among traits causes the unequal distribution of genetic variance among multi-trait combinations, and ultimately can cause the number of eigenvectors with statistically significant variance (known as the matrix rank) to be less than the number of measured traits. Differences in rank may arise even when the magnitude of individual pairwise trait covariances do not significantly differ between **G_I_** and **G_O_**. To assess differences in rank, we performed factor analytic modelling [45, 46] on each dataset. We performed likelihood ratio tests, comparing models where **G** is estimated at full rank, then sequentially at reduced rank.

Even when both full rank, matrices may differ in shape. If, for example, non-additive effects tended to strengthen genetic correlations, then the leading eigenvalues of **G_I_** would explain a higher proportion of variance than the leading eigenvalues of **G_O_** (*e.g*., blue *versus* red **G** in **Figure 4B**). To compare the distribution of eigenvalues, we performed eigenanalyses of **G_O_** and **G_I_** using the *eigen* function in base R [42]. To place confidence intervals on the eigenvalues, we projected the eigenvectors of the REML estimates of **G_O_** and **G_I_** through the 10,000 REML-MVN samples of the respective matrix using the equation [47]:

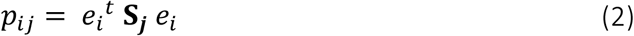

where *p_ij_* is the estimate of variance in the *i*th eigenvector (*e_i_*; *i* = 1 to 11), in the *j*th REML-MVN sample (**S_*j*_**; *j* = 1 to 10,000), and *t* indicates the transpose. We tested for significant differences in shape between **G_O_** and **G_I_** by comparing the 95% CI of the variance associated with each eigenvector.

Third, we evaluated the effect of inbreeding on matrix orientation (**Figure 4C**). We calculated the dot product between the 11 eigenvectors of **G_O_** and **G_I_**. The absolute value of a dot product of two normalised vectors ranges from zero to one, with one indicating that the two eigenvectors represent the same multi-trait combination, and zero that the two multi-trait combinations are orthogonal. Additionally, we compared the two **G** in the space of the outbred **G** by projecting (**Equation 2**) the eigenvectors of the REML estimate of **G_O_** through the REML estimates of **G_O_** and **G_I_** and through the 10,000 REML-MVN samples of **G_O_** and **G_I_**. To test whether *V_A_* differed in any of the 11 eigenvectors, we paired REML-MVN samples and calculated the ratio of variances (*V_A_O__/V_A_I__*) for each eigenvector, with statistical significance evaluated by comparing the 95% CI against 1.0. This projection allows us to test whether dimensions with relatively more (less) variance in **G_O_** also have more (less) variance in **G_I_**, and, therefore, whether the additive model adequately accounts for the effect of inbreeding across multivariate traits associated with different amounts of *V_A_*.

While eigenvectors of **G** are informative of the distribution of *V_A_*, and allow us to specifically probe whether certain eigenvectors depart from additive expectations, there is no inherent constraint for changes in *V_A_* (or evolution) to align with these constructs. Therefore, in our final analysis we used an eigentensor approach to explicitly identify the multi-trait combinations that most strongly departed from the expectation under additive gene action, irrespective of the distribution of variation in the traits. The eigentensor:

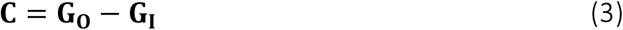

describes differences between the **G** on the absolute scale [48], which is most relevant for predicting impacts of the genetic difference on evolution [48, 49]. The largest absolute eigenvalue of **C** describes the multi-trait combination where the absolute difference in *V_A_* is greatest between **G_O_** and **G_I_**, and hence the multi-trait combination associated with the greatest departure from the additive model, and where evolutionary outcomes might become least predictable from the outbred **G**. To calculate confidence intervals, we paired the 10,000 REML-MVN sample matrices of **G_O_** and of **G_I_** and calculated **C** for each of the 10,000 pairs. We calculated confidence intervals for the eigenvalues of **C** by projecting (**Equation 2**) the eigenvectors of the REML point estimate of **C** through the 10,000 REML-MVN samples of **C**. Multi-trait combinations deviating from additive expectations were identified where the 95% CI of the eigenvalue of **C** did not include zero.

## Results

We first determine how inbreeding affected individual trait means and variances. We then describe the multivariate genetic variation in wing traits in our outbred population of *D. serrata*, and, finally, present evidence of how inbreeding influenced that multivariate genetic variation.

### The effect of inbreeding on individual traits

Inbreeding scarcely affected wing trait means (**Figure 5A**; **Table S1**). The maximum ID was just 3.28%, with the mean ID <1% across the 11 traits. The 95% CI of the 3,502 estimates spanned ranges of >10%, and, notably, this range was particularly large for ILD1.9 (**Figure 5A**). That is, the difference in trait value between the inbred and outbred offspring of a sire could be quite substantial, but there was no consistency among sires in the direction of the deviation. These results support no consistent, directional dominance of effects, thereby allowing us to test the (1 + *F*)*V_A_* prediction (**Figure 1**).

**Figure 5.**
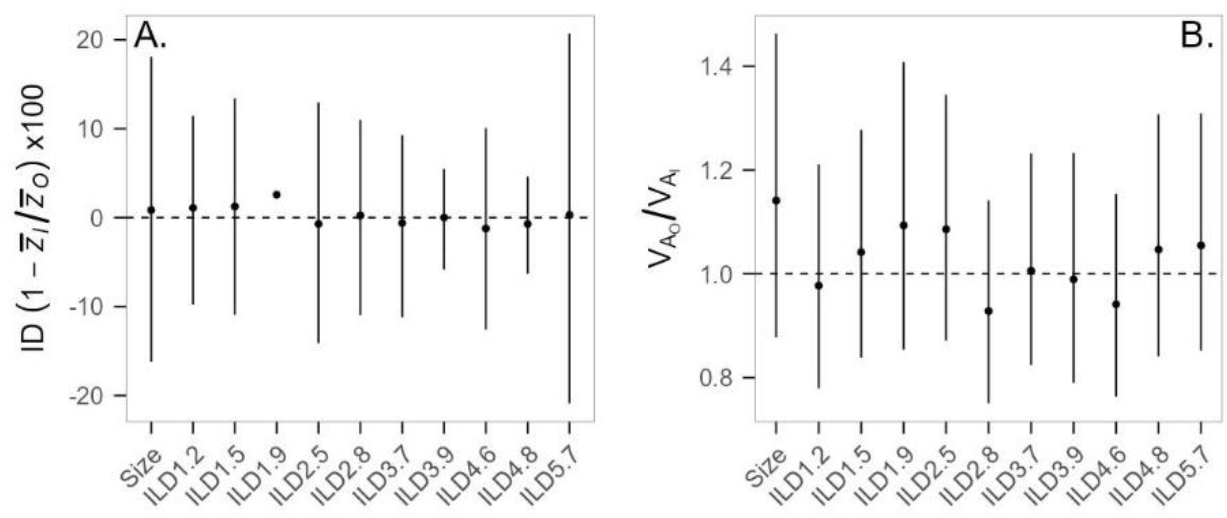
The effect of inbreeding on trait means (**A**) and additive genetic variances (**B**) for 11 wing traits. **A**. Inbreeding depression (ID; ±95% CI) is presented as the percentage change in trait mean with inbreeding, assuming individuals are fully inbred (see methods). The horizonal dashed line at zero represents no ID, while positive (negative) values indicate inbreeding increased (decreased) trait mean. CI for ILD1.9 were large (−47.78; 50.47) and are omitted to improve visualisation of CI for other traits. **B**. The ratio of additive genetic variance without (*V_A_O__*) and with (*V_A_I__*) inbreeding. The horizontal dashed line represents *V_A_O__* = *V_A_I__*, with values above (below) the line having less (more) *V_A_* with inbreeding than expected if effects are additive.

We found statistical support for *V_A_* in all 11 individual traits in the outbred (**Table S2**) and inbred (**Table S3**) populations. For each trait, the magnitude of *V_A_* was similar (**Tables S2** and **S3**), and 95% CI of the ratio of variances substantially overlapped 1.0 (**Figure 5B**). Thus, there was no evidence that inbreeding affected the *V_A_* of any trait more (or less) than predicted by additive effects. Similarly, the 95% CI for the ratio of the total *V_A_* overlapped one (median: 1.016; 95% CI: 0.922-1.120).

### Genetic (co)variance in wing traits

In the absence of inbreeding, absolute genetic correlations ranged from 0.01 to 0.76, with a median of 0.22, and slightly more (56%) negative than positive estimates (**Table S2**). Correlations between size and shape traits, with a median absolute value of 0.11, were generally weaker than those among shape traits (**Table S2**). We found weak statistical support for genetic variance in all 11 dimensions of **G_O_** (comparison of rank of 11 *versus* 10: *χ^2^* = 3.66; d.f. = 1; *P* = 0.056; see **Table S4**). Collectively, these observations (individual trait variance, modest pairwise correlations and high rank of **G_O_**) suggest that genetic variation was present across the phenotypic trait space. However, the eigen-analysis revealed the uneven distribution of this variation. The first eigenvector of **G_O_** explained 34.47% of the variance and, notably, the last four eigenvectors of **G_O_** had very low *V_A_*, each accounting for <1% of the total genetic variance (**Figure 6A**; **Table S4**).

**Figure 6.**
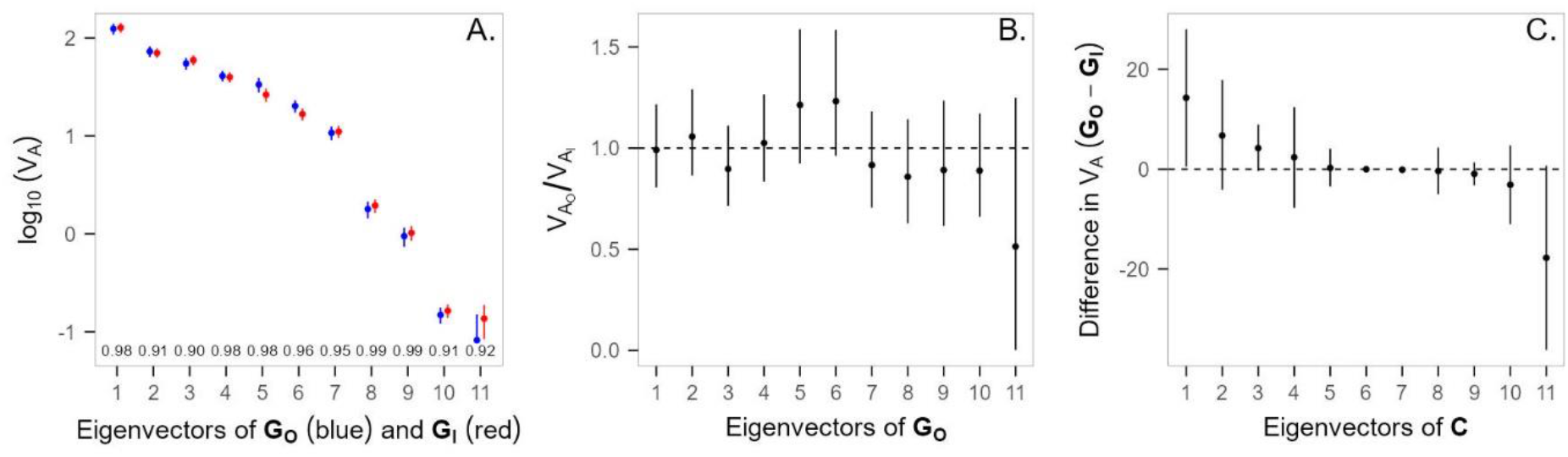
The effect of inbreeding on **G**. **A**. The distribution of eigenvalues (±90% CI) of **G_O_** (blue) and **G_I_** (red) are plotted on the log scale. Dot products between eigenvectors are shown along the x-axis. The lower CI for the smallest eigenvalue of **G_O_** is not shown, but is 0.35×10^−4^ (−3.5 on the log scale). **B.** The ratio (±95% CI) of additive genetic variance (*V_A_*) in **G_O_** (*V_A_O__*) and **G_I_** (*V_A_I__*) in the eigenvectors of **G_O_**. The horizontal dashed line represents the same magnitude of variance in each matrix for a given eigenvector, with values above (below) the line having deflation (inflation) of *V_A_* relative to that expected by inbreeding when effects are additive. **C.** Eigenvalues (±95% CI) of the difference matrix, **C**. Eigenvalues above (below) the zero (dashed line) indicates deflation (inflation) of variance by inbreeding, after accounting for the additive expectation.

### Does the effect of inbreeding on genetic (co)variance deviate from the additive predictions?

Individual pairwise genetic trait covariances were similar (overlapping 95% CI) with and without inbreeding, although four pairwise correlations were only statistically supported as non-zero in one dataset (three in **G_I_** only, and one in **G_O_** only), with two of these estimates of opposing sign between the inbred and outbred estimates (**Tables S2** and **S3**). We found strong statistical support for all 11 dimensions of **G_I_** (comparison of rank of 11 *versus* 10: *χ*^2^ = 17.28; d.f. = 1; *P* < 0.0001; **Table S5**). Although statistical support for full rank was stronger for **G_I_** than for **G_O_**, we suspect this reflects slightly greater power in the former analysis, rather than reflecting differences in genetic variation in the final dimension of each **G** (**Figure 6A; Tables S3** and **S4**).

**G_O_** and **G_I_** were also markedly similar in terms of their shape (**Figure 4B**) and orientation (**Figure 4C**). There was substantial overlap of 95% CI for each of the 11 eigenvalues, and all dot products exceeded 0.90, suggesting that the same trait combinations were associated with similar variance in each matrix (**Figure 6A**; **Tables S4** and **S5**). This interpretation is supported by the observation that, for all eigenvectors of **G_O_**, the 95% CI for the ratio of the variances substantially overlapped one (**Figure 6B**), demonstrating that the magnitude of *V_A_* was similar for **G_I_** and **G_O_** across the eigenvectors of **G_O_**. In particular, the four phenotypic dimensions that had the lowest genetic variance in **G_O_** were similarly depauperate of genetic variance in **G_I_**. While there was approximately twice as much variance in **G_I_** (compared to **G_O_**) for the last eigenvector of **G_O_** (**Figure 6B**), the 95% CI overlapped zero, and the slightly greater power in the inbred analysis would result in less downward bias in this eigenvalue [50, 51].

Finally, the eigentensor supported strong similarity of **G_O_** and **G_I_**. The absolute eigenvalues for *c*_2_-*c*_10_ were all very small and had 95% CI that overlapped zero (**Figure 6C**; **Table S5**). The remaining two eigenvectors of **C** (*c*_1_ and *c*_*G*_11__) are the most important in that they reflect the two multi-trait combinations that differ most between **G_O_** and **G_I_**. While opposite in sign (reflecting inbreeding decreasing or increasing genetic variance relative to the expectation under only additive effects) *c*_1_and *c*_11_ are comparable in absolute magnitude and are each associated with the plane defined by the first few eigenvectors of **G** (capturing approximately 20% of the variance in each **G**; compare the eigenvectors of **G** in **Tables S4** and **S5** to *c*_1_ and *c*_11_ in **Table S6**). Thus, the data are suggestive of a subtle change in the shape and/or orientation associated with regions of trait space with substantial *V_A_*. Sampling error has been demonstrated to inflate leading eigenvalues, conflating biological and statistical signals [50, 51]. Here, independent sampling error associated with each **G** may contribute to apparent differences between them. The CI associated with both *c*_1_ and *c*_11_ are relatively large (**Figure 6C**), which we suggest is expected if the differences between the **G** are due to an independent realisation of **G** (*i.e*., sampling error, which is incorporated into the eigentensor), rather than an effect of inbreeding. Therefore, we interpret the eigenvalues of **C** cautiously, and suggest that the eigentensor supports our previous conclusions of striking similarity between the two **G**.

## Discussion

The maintenance of high fitness of populations ultimately depends on the presence of additive genetic variation (*V_A_*) and its underlying genetic architecture. Loci with rare, recessive, alleles may contribute substantially to *V_A_*, but the rarity of such variants is a major challenge to characterising their genetic effects and contributions. Here, we provide an innovative test of genetic architecture underlying *Drosophila* wing traits by controlling for sampling effects and increasing the frequency of homozygotes at loci currently segregating in the population. We found evidence that the effects of inbreeding on the additive genetic (co)variances (**G**) were consistent with expectations when contributing variants have additive (not dominant) effects. Thus, our results suggest that the average dominance coefficient is small compared to the average additive effect and/or variants affecting the traits are at intermediate frequencies (*i.e*., *d* ≪ *a* and/or *q* ~ *p*; see **Figure 1**).

For our specific trait set, we expected **G** to be impacted by homozygosity for several reasons. First, while genetic effects do not map simply to genetic variance components [6], detection of dominance genetic variance for *D. serrata* wing traits [30] suggested the presence of loci with recessive alleles. In *D. melanogaster*, wing traits are affected by new mutations with an average dominance coefficient of 0.25 [52], and mapping studies have implicated (partially) recessive candidate loci [53], while analyses of wild-caught flies implicated a recessive allele at one of the best-supported candidate causal variants (Epidermal growth factor receptor, *Egfr*) [54]. Second, some dimensions of wing trait space of *D. serrata* [this study, and 29, 30] and *D. melanogaster* [40] have very low *V_A_*, which is expected to reflect the predominant contribution from rare alleles [25]; multivariate GWAS of wing shape in *D. melanogaster* has implicated causal loci with rare alleles [55]. Finally, pleiotropic alleles jointly affect *Drosophila* wings and fitness [56–58], and loci with such effects are predicted to contribute to increased variance under inbreeding due to dominance effects [16].

Despite these reasons to expect dominant gene action to contribute to *V_A_*, we found that **G_I_** did not depart from the additive prediction. We were able to robustly estimate and compare all phenotypic dimensions of **G** including dimensions with very low *V_A_*, which define multi-trait combinations that are predicted to be enriched for rare, recessive alleles. Despite this power, the contribution of rare, recessive alleles to *V_A_* was undetectable against a background of additive effects and/or intermediate-frequency alleles. Indeed, manipulative laboratory experiments have demonstrated polygenic architecture with additive effects and common alleles for wing shape traits in *D. melanogaster* [59, 60], which is consistent with rapid evolution of wings in the wild [61]. Notably, other studies that have attempted to directly test the contribution of (partially) recessive, rare alleles to variation in a range of different quantitative traits (including fitness-related traits) have, to date, found little evidence for a major contribution [62–64].

Although relatively small dominance coefficients or similar frequencies of the alternative alleles at a locus are plausible explanations for our results, we consider several possible alternative explanations. First, inflation of *V_A_* due to non-additive genetic effects is predicted to be maximised when the inbreeding coefficient, *F*, is 0.4 [65], considerably above that achieved here via one generation of full-sib inbreeding (*F*~0.25). We aimed to exclude any contribution from divergence in allele frequency between the inbred and outbred populations due to chance sampling (drift), precluding multi-generational inbreeding from our design. Increasing *F* increases the frequency of homozygotes (in terms of both number of sampled individuals for a given locus, and the number of loci), therefore also increasing the power to detect an effect, and we note that deviation of **G_I_** from the additive model may have been detectable with larger *F*.

Second, while our experimental design was expected to result in >25 times higher per-locus homozygosity for rare (*q* < 0.01) alleles in the inbred pedigree (**Figure 3**), if these alleles were rare due to highly deleterious effects on pre-adult fitness, viability selection may have influenced the estimate of **G_I_**, but not **G_O_**. That is, despite the presence of segregating rare recessive allele at a locus, all (inbred and outbred) siblings shared the same phenotype because the rare allele homozygote was not sampled. Such viability selection in the inbred population, decreasing *V_A_*, would decrease **G_I_** below (1 + *F*)*V_A_*. Therefore, a compensatory inflation of **G_I_** (through the contribution of non-lethal rare recessive alleles at other loci) would be necessary to explain our observation of **G_O_** = **G_I_**.

If viability selection indeed precluded individuals from being sampled for their wing phenotypes, then wing shape/size could not be the direct cause of this fitness effect. Nonetheless, such selection can have consequences for wing variation via pleiotropic effects, as previously demonstrated for *D. serrata* wing shape and size [30, 56, 57]. The joint distribution of pleiotropic allelic effects on fitness and other traits remains logistically difficult to examine, but, for traits under stabilising selection, additive effects are predicted to be correlated, such that alleles causing large (small) changes in trait value also cause large (small) decreases in fitness [66]. However, theoretical studies have shown that dominance effects on fitness can emerge when the allelic effects on traits contributing to fitness are additive [67, 68], suggesting that, in contrast to additive pleiotropic effects, the magnitude of the dominance coefficients of pleiotropic alleles may be uncorrelated between trait and fitness. If this is the case, then loci at which *q* → 0 due to selection may typically have *d*~0 for morphological traits but *d* ≫ 0 for fitness. Notably, if inbreeding alters *V_A_* for fitness and the rate of evolution, the evolution of wing *V_A_*, and the distribution of size and wing shape, will also be affected.

Considerable attention has been given to how non-additive gene action may affect *V_A_* following population bottlenecks, where the distribution of allele frequencies is altered by genetic drift, including fixation at some loci, increasing homozygosity [69]. Here, we were focused on the contributions of dominance effects under the allele frequency spectra that had evolved in the population – specifically, the contribution of rare alleles, which are readily lost through bottlenecks [70]. If rare alleles contribute more to multi-trait combinations with low *V_A_*, then random genetic drift under population decline might differentially impact *V_A_* across multivariate trait space. Notably, reduction to a population size of one male and one female for one generation did not, on average, change the shape or orientation of **G** for *D. melanogaster* wings [26]. This observation suggests the loci that most rapidly lost variation under drift contributed similarly across the 6-dimensional trait space, inconsistent with the expectation that dimensions with low *V_A_* are enriched for rare alleles. However, estimating 52 **G** necessitated relatively small sample sizes for each [26], and estimation error may have inhibited detection of shape or orientation change.

Finally, we stress that we have not dealt with *dominance variance*, for which we had low power to estimate (*e.g*., only 16 double-first cousin families), particularly in the presence of inbreeding [3]. Our focus on **G** and *dominance effects* reflects their immediate importance to evolution. Inbreeding, which is inevitable in small, fragmented populations, may cause non-proportional changes to **G** if genetic architecture varies across multivariate trait space. Such a non-proportional change may alter evolutionary trajectories and/or the capacity of populations to purge mutations, which is critical for the maintenance of population fitness [71]. While our results suggest no deviation from the additive prediction, this may not be true of other trait sets, or in other taxa. Greater contribution of non-additive gene action to *V_A_* of fitness than morphological traits has been posited, although empirical evidence for a difference is equivocal [69, 72, 73]. Indeed, historically applied approaches (changes in trait mean with inbreeding, or estimates of non-additive variance components) can only reveal non-additive gene action under some conditions, while studies focusing on changes in the multivariate distribution of *V_A_* may provide novel insights into the prevalence and evolutionary relevance of non-additive gene action. Experiments that can characterise the phenotypic effects of rare alleles are extremely challenging, but extending our understanding of gene action at such loci, and the distribution of pleiotropic effects on phenotypic traits and fitness, is critical to improving our ability to predict population evolutionary responses to rapid environment change and population decline.

## Acknowledgements

We thank S. Chenoweth for providing the base population, and A. Reddiex, D. Sun, C. Liddington, H. North, J. Price, R. Yong, W. Arnold, C. Conradsen, N. Appleton, A. Oliveros-Sandino, G. Hayes, and A. Kannan for technical assistance. This manuscript was improved by feedback from anonymous reviewers and J. Hadfield. This work was funded by the Australian Research Council and The University of Queensland.

