## Supplementary material for "Using inbreeding to test the contribution of non-additive genetic effects to additive genetic variance: a case study in *Drosophila serrata*": Table S1

**Table S1** | Estimates of inbreeding depression (ID) for 11 wing traits. Raw phenotypic means for outbred ($\bar{z}_O$) and inbred ($\bar{z}_{Iraw}$) flies are shown, together with the inbred mean calculated from linear extrapolation assuming individuals are completely inbred ($\bar{z}_I$). ID was calculated as: $1-(\bar{z}_I/\bar{z}_O)$.

| Trait | $\bar{z}_O$ | $\bar{z}_{Iraw}$ | $\bar{z}_I$ | ID |
|---|---|---|---|---|
| Wing size | 331.72 | 330.95 | 328.64 | 0.0093 |
| ILD1.2 | 700.53 | 698.68 | 693.13 | 0.0106 |
| ILD1.5 | 560.23 | 558.47 | 553.19 | 0.0126 |
| ILD1.9 | 133.31 | 132.35 | 129.47 | 0.0288 |
| ILD2.5 | 524.07 | 525.00 | 527.79 | -0.0071 |
| ILD2.8 | 589.54 | 589.92 | 591.06 | -0.0026 |
| ILD3.7 | 586.02 | 585.16 | 582.58 | 0.0059 |
| ILD3.9 | 770.94 | 770.76 | 770.22 | 0.0009 |
| ILD4.6 | 545.44 | 547.12 | 552.16 | -0.0123 |
| ILD4.8 | 734.13 | 735.36 | 739.05 | -0.0067 |
| ILD5.7 | 260.81 | 261.03 | 261.69 | -0.0034 |

**Table S2** | Additive genetic covariance matrix for the outbred data set ($\mathbf{G_O}$). Additive genetic variances (90% CI) are on the diagonals, covariances (95% CI) are below the diagonals, and correlations (95% CI) above the diagonals.

| | Size | ILD1.2 | ILD1.5 | ILD1.9 | ILD2.5 | ILD2.8 | ILD3.7 | ILD3.9 | ILD4.6 | ILD4.8 | ILD5.7 |
|---|---|---|---|---|---|---|---|---|---|---|---|
| Size | **23.68**<br>*19.80; 27.57* | 0.06<br>*-0.07; 0.18* | -0.08<br>*-0.20; 0.05* | -0.18<br>*-0.32; -0.05* | -0.30<br>*-0.41; -0.18* | 0.17<br>*0.04; 0.28* | 0.04<br>*-0.08; 0.16* | 0.15<br>*0.02; 0.27* | 0.06<br>*-0.07; 0.18* | 0.29<br>*0.17; 0.41* | 0.01<br>*-0.12; 0.14* |
| ILD1.2 | 2.21<br>*-2.29; 6.63* | **52.04**<br>*44.82; 59.09* | 0.31<br>*0.20; 0.41* | 0.37<br>*0.26; 0.47* | -0.05<br>*-0.17; 0.06* | 0.75<br>*0.69; 0.80* | -0.38<br>*-0.47; -0.28* | -0.44<br>*-0.53; -0.33* | -0.41<br>*-0.50; -0.31* | -0.52<br>*-0.60; -0.43* | -0.10<br>*-0.21; 0.02* |
| ILD1.5 | -2.45<br>*-6.43; 1.51* | 14.63<br>*9.19; 20.03* | **42.80**<br>*37.24; 48.35* | 0.25<br>*0.14; 0.36* | -0.23<br>*-0.34; -0.12* | 0.14<br>*0.03; 0.25* | -0.57<br>*-0.64; -0.49* | -0.34<br>*-0.44; -0.24* | -0.53<br>*-0.61; -0.45* | -0.08<br>*-0.19; 0.04* | 0.47<br>*0.38; 0.55* |
| ILD1.9 | -4.57<br>*-8.00; -1.17* | 14.04<br>*9.13; 19.00* | 8.63<br>*4.45; 12.80* | **26.99**<br>*22.86; 31.01* | 0.26<br>*0.14; 0.38* | -0.33<br>*-0.43; -0.22* | -0.17<br>*-0.27; -0.05* | -0.67<br>*-0.73; -0.59* | -0.02<br>*-0.13; 0.11* | -0.68<br>*-0.75; -0.60* | -0.20<br>*-0.32; -0.08* |
| ILD2.5 | -9.67<br>*-14.02; -5.32* | -2.63<br>*-8.14; 2.85* | -10.21<br>*-15.03; -5.12* | 9.21<br>*4.94; 13.44* | **44.90**<br>*38.76; 50.88* | -0.18<br>*-0.29; -0.06* | -0.03<br>*-0.13; 0.08* | -0.57<br>*-0.64; -0.48* | 0.09<br>*-0.02; 0.21* | -0.58<br>*-0.65; -0.49* | 0.02<br>*-0.09; 0.13* |
| ILD2.8 | 5.16<br>*1.24; 9.03* | 34.60<br>*28.30; 40.85* | 6.01<br>*1.38; 10.60* | -10.93<br>*-15.14; -6.79* | -7.58<br>*-12.45; -2.67* | **41.30**<br>*35.85; 46.64* | -0.29<br>*-0.39; -0.19* | -0.01<br>*-0.13; 0.10* | -0.41<br>*-0.50; -0.32* | -0.08<br>*-0.19; 0.04* | 0.05<br>*-0.06; 0.17* |
| ILD3.7 | 1.30<br>*-2.40; 4.93* | -17.03<br>*-22.32; -11.78* | -23.31<br>*-28.19; -18.38* | -5.36<br>*-9.06; -1.48* | -1.18<br>*-5.63; 3.41* | -11.47<br>*-15.90; -7.04* | **38.53**<br>*33.77; 43.36* | 0.22<br>*0.11; 0.33* | 0.76<br>*0.71; 0.81* | -0.06<br>*-0.17; 0.05* | 0.23<br>*0.13; 0.33* |
| ILD3.9 | 2.61<br>*0.28; 4.90* | -11.60<br>*-14.99; -8.19* | -8.25<br>*-11.13; -5.33* | -12.81<br>*-15.56; -9.93* | -14.10<br>*-17.33; -10.79* | -0.31<br>*-3.00; 2.42* | 5.11<br>*2.44; 7.68* | **13.66**<br>*11.74; 15.54* | -0.07<br>*-0.19; 0.04* | 0.64<br>*0.56; 0.71* | -0.24<br>*-0.35; -0.13* |
| ILD4.6 | 1.80<br>*-1.97; 5.47* | -18.24<br>*-23.74; -12.76* | -21.52<br>*-26.38; -16.55* | -0.49<br>*-4.17; 3.37* | 3.74<br>*-0.90; 8.32* | -16.35<br>*-20.99; -11.70* | 29.06<br>*24.22; 34.06* | -1.70<br>*-4.36; 0.90* | **38.04**<br>*33.20; 42.97* | 0.04<br>*-0.08; 0.15* | 0.10<br>*-0.01; 0.22* |
| ILD4.8 | 5.20<br>*2.92; 7.55* | -13.90<br>*-17.49; -10.39* | -1.94<br>*-4.73; 0.83* | -13.07<br>*-15.78; -10.36* | -14.36<br>*-17.62; -11.09* | -1.82<br>*-4.53; 0.85* | -1.32<br>*-3.87; 1.17* | 8.77<br>*6.92; 10.64* | 0.90<br>*-1.70; 3.53* | **13.79**<br>*11.96; 15.70* | -0.05<br>*-0.17; 0.06* |
| ILD5.7 | 0.13<br>*-2.94; 3.18* | -3.52<br>*-7.65; 0.62* | 15.29<br>*11.46; 19.27* | -5.24<br>*-8.45; -2.02* | 0.82<br>*-3.00; 4.60* | 1.69<br>*-1.92; 5.26* | 7.09<br>*3.69; 10.63* | -4.42<br>*-6.60; -2.23* | 3.21<br>*-0.25; 6.66* | -0.98<br>*-3.14; 1.14* | **24.66**<br>*21.52; 27.97* |

**Table S3** | Additive genetic covariance matrix for the inbred data set (**G$_I$**). Additive genetic variances (90% CI) are on the diagonals, covariances (95% CI) are below the diagonals, and correlations (95% CI) above the diagonals.

| | Size | ILD1.2 | ILD1.5 | ILD1.9 | ILD2.5 | ILD2.8 | ILD3.7 | ILD3.9 | ILD4.6 | ILD4.8 | ILD5.7 |
|---|---|---|---|---|---|---|---|---|---|---|---|
| Size | **20.75**<br>*17.94; 23.54* | 0.10<br>*-0.01; 0.21* | -0.09<br>*-0.20; 0.02* | -0.20<br>*-0.32; -0.09* | -0.28<br>*-0.38; -0.18* | 0.20<br>*0.09; 0.30* | 0.08<br>*-0.03; 0.19* | 0.16<br>*0.05; 0.26* | -0.02<br>*-0.12; 0.09* | 0.21<br>*0.11; 0.32* | 0.04<br>*-0.07; 0.15* |
| ILD1.2 | 3.40<br>*-0.19; 7.06* | **53.26**<br>*46.75; 59.76* | 0.22<br>*0.13; 0.32* | 0.32<br>*0.22; 0.41* | 0.14<br>*0.04; 0.24* | 0.79<br>*0.75; 0.83* | -0.39<br>*-0.47; -0.30* | -0.46<br>*-0.53; -0.37* | -0.45<br>*-0.53; -0.36* | -0.60<br>*-0.66; -0.53* | -0.06<br>*-0.16; 0.04* |
| ILD1.5 | -2.70<br>*-5.83; 0.45* | 10.52<br>*5.69; 15.37* | **41.09**<br>*36.21; 46.07* | 0.21<br>*0.10; 0.31* | -0.23<br>*-0.33; -0.13* | 0.09<br>*-0.01; 0.19* | -0.57<br>*-0.64; -0.50* | -0.29<br>*-0.38; -0.19* | -0.53<br>*-0.60; -0.45* | -0.03<br>*-0.13; 0.08* | 0.49<br>*0.41; 0.56* |
| ILD1.9 | -4.59<br>*-7.16; -1.99* | 11.70<br>*7.30; 15.91* | 6.65<br>*2.99; 10.22* | **24.69**<br>*21.20; 28.14* | 0.35<br>*0.24; 0.45* | -0.31<br>*-0.41; -0.21* | -0.17<br>*-0.27; -0.06* | -0.69<br>*-0.75; -0.63* | -0.03<br>*-0.13; 0.08* | -0.61<br>*-0.68; -0.54* | -0.19<br>*-0.30; -0.08* |
| ILD2.5 | -8.17<br>*-11.32; -4.98* | 6.56<br>*1.79; 11.26* | -9.56<br>*-13.81; -5.27* | 11.15<br>*7.57; 14.60* | **41.35**<br>*36.30; 46.23* | -0.03<br>*-0.13; 0.07* | -0.08<br>*-0.18; 0.01* | -0.58<br>*-0.65; -0.51* | 0.00<br>*-0.09; 0.10* | -0.67<br>*-0.73; -0.61* | -0.08<br>*-0.19; 0.02* |
| ILD2.8 | 5.98<br>*2.73; 9.25* | 38.49<br>*32.47; 44.62* | 3.71<br>*-0.46; 7.90* | -10.32<br>*-14.08; -6.60* | -1.18<br>*-5.48; 3.03* | **44.49**<br>*39.32; 49.78* | -0.28<br>*-0.37; -0.19* | -0.05<br>*-0.15; 0.05* | -0.42<br>*-0.51; -0.34* | -0.24<br>*-0.33; -0.14* | 0.06<br>*-0.04; 0.17* |
| ILD3.7 | 2.25<br>*-0.76; 5.18* | -17.60<br>*-22.40; -12.72* | -22.79<br>*-27.29; -18.41* | -5.08<br>*-8.34; -1.84* | -3.36<br>*-7.29; 0.58* | -11.61<br>*-15.78; -7.40* | **38.30**<br>*33.99; 42.58* | 0.20<br>*0.10; 0.29* | 0.77<br>*0.72; 0.81* | 0.03<br>*-0.07; 0.13* | 0.24<br>*0.14; 0.33* |
| ILD3.9 | 2.67<br>*0.83; 4.47* | -12.39<br>*-15.38; -9.34* | -6.88<br>*-9.39; -4.35* | -12.78<br>*-15.18; -10.32* | -13.90<br>*-16.59; -11.11* | -1.25<br>*-3.67; 1.20* | 4.52<br>*2.24; 6.82* | **13.80**<br>*12.14; 15.47* | -0.07<br>*-0.17; 0.03* | 0.61<br>*0.54; 0.67* | -0.22<br>*-0.32; -0.12* |
| ILD4.6 | -0.48<br>*-3.63; 2.58* | -20.72<br>*-25.78; -15.75* | -21.51<br>*-26.02; -17.05* | -0.81<br>*-4.05; 2.55* | 0.16<br>*-3.88; 4.20* | -17.98<br>*-22.34; -13.69* | 30.15<br>*25.40; 34.78* | -1.61<br>*-3.91; 0.69* | **40.41**<br>*35.84; 45.01* | 0.15<br>*0.05; 0.25* | 0.11<br>*0.01; 0.21* |
| ILD4.8 | 3.54<br>*1.73; 5.33* | -15.83<br>*-18.91; -12.64* | -0.58<br>*-2.99; 1.84* | -11.04<br>*-13.35; -8.76* | -15.72<br>*-18.55; -12.86* | -5.76<br>*-8.20; -3.31* | 0.61<br>*-1.62; 2.83* | 8.21<br>*6.63; 9.76* | 3.54<br>*1.15; 5.91* | **13.18**<br>*11.53; 14.79* | 0.00<br>*-0.11; 0.11* |
| ILD5.7 | 0.91<br>*-1.54; 3.28* | -2.28<br>*-5.90; 1.34* | 15.08<br>*11.72; 18.54* | -4.65<br>*-7.39; -1.88* | -2.57<br>*-5.69; 0.52* | 2.05<br>*-1.13; 5.30* | 7.13<br>*4.06; 10.11* | -4.01<br>*-5.89; -2.14* | 3.53<br>*0.39; 6.54* | -0.02<br>*-1.85; 1.81* | **23.39**<br>*20.57; 26.19* |

**Table S4** | Eigen-analysis of $\mathbf{G_O}$. Eigenvalues (λ) are bolded, with CI in italics; eigenvectors (normalised trait loadings) are in the lower part of the table. Factor analysis showed weak statistical support for variance in $e_{11}$ ($\chi^2$ = 3.66; d.f. = 1; *P* = 0.056). The REML estimation in WOMBAT is constrained to be positive-definite (no negative eigenvalues), precluding transformation to the Tracy-Widom distribution; the factor analytic approach is as robust as the TW transformation against false positives, but may be more conservative for smaller eigenvalues (see Sztepanacz and Blows, 2017). Here, we rely on the factor analytic results with respect to the rank of the matrix, and report only the 95% CI for comparison to the eigenvalues of $\mathbf{G_I}$, presented in **Table S4**.

| | $e_1$ | $e_2$ | $e_3$ | $e_4$ | $e_5$ | $e_6$ | $e_7$ | $e_8$ | $e_9$ | $e_{10}$ | $e_{11}$ |
|---|---|---|---|---|---|---|---|---|---|---|---|
| λ | **124.24** | **72.82** | **54.94** | **41.01** | **33.43** | **20.23** | **10.75** | **1.79** | **0.95** | **0.15** | **0.08** |
| 95% CI | *105.29; 142.88* | *61.84; 84.08* | *45.77; 64.07* | *34.90; 47.35* | *26.63; 40.16* | *16.77; 23.78* | *8.73; 12.82* | *1.37; 2.20* | *0.69; 1.20* | *0.12; 0.18* | *<0.00; 0.16* |
| Size | 0.00 | -0.24 | -0.15 | 0.09 | -0.16 | 0.90 | -0.25 | -0.10 | 0.02 | 0.00 | 0.00 |
| ILD1.2 | -0.54 | 0.12 | -0.47 | 0.17 | -0.22 | -0.07 | 0.02 | 0.18 | -0.20 | -0.29 | 0.49 |
| ILD1.5 | -0.40 | -0.02 | 0.55 | 0.40 | 0.00 | 0.01 | 0.07 | -0.45 | 0.32 | -0.26 | 0.09 |
| ILD1.9 | -0.11 | 0.44 | 0.08 | 0.08 | -0.55 | 0.02 | -0.13 | 0.18 | -0.05 | -0.18 | -0.62 |
| ILD2.5 | 0.07 | 0.65 | -0.08 | -0.21 | 0.54 | 0.25 | -0.04 | -0.10 | 0.08 | -0.38 | 0.03 |
| ILD2.8 | -0.39 | -0.24 | -0.49 | 0.07 | 0.34 | -0.07 | 0.16 | -0.16 | 0.21 | 0.04 | -0.57 |
| ILD3.7 | 0.42 | -0.02 | -0.31 | 0.42 | -0.04 | -0.24 | -0.46 | 0.02 | 0.49 | -0.18 | 0.07 |
| ILD3.9 | 0.11 | -0.34 | 0.00 | -0.20 | 0.00 | -0.19 | -0.31 | -0.37 | -0.49 | -0.55 | -0.14 |
| ILD4.6 | 0.43 | 0.10 | -0.23 | 0.36 | -0.17 | 0.08 | 0.66 | -0.31 | -0.20 | -0.11 | -0.02 |
| ILD4.8 | 0.09 | -0.35 | 0.14 | -0.12 | 0.03 | 0.11 | 0.35 | 0.56 | 0.27 | -0.56 | -0.05 |
| ILD5.7 | 0.00 | -0.02 | 0.18 | 0.62 | 0.43 | 0.06 | -0.13 | 0.37 | -0.46 | 0.04 | -0.13 |

**Table S5** | Eigen-analysis of $\mathbf{G_I}$. Eigenvalues (λ) are bolded with CI in italics; eigenvectors (normalised trait loadings) are in the lower part of the table. Factor analysis showed strong statistical support for variance in $e_{11}$ ($\chi^2$ = 17.28; d.f. = 1; $P$ < 0.0001). 95% CI are for comparison to the eigenvalues of $\mathbf{G_O}$, presented in **Table S2**. See Table S2 for further information.

| | $e_1$ | $e_2$ | $e_3$ | $e_4$ | $e_5$ | $e_6$ | $e_7$ | $e_8$ | $e_9$ | $e_{10}$ | $e_{11}$ |
|---|---|---|---|---|---|---|---|---|---|---|---|
| λ | **127.75** | **70.29** | **59.45** | **39.82** | **26.35** | **16.70** | **11.08** | **1.95** | **1.02** | **0.16** | **0.14** |
| 95% CI | *110.13; 145.31* | *61.35; 79.40* | *50.80; 68.08* | *34.29; 45.34* | *21.45; 31.38* | *13.85; 19.56* | *9.25; 12.97* | *1.57; 2.31* | *0.81; 1.24* | *0.13; 0.20* | *0.08; 0.20* |
| Size | -0.01 | -0.16 | -0.24 | 0.05 | -0.23 | 0.92 | 0.00 | -0.13 | 0.02 | 0.00 | -0.01 |
| ILD1.2 | -0.56 | 0.21 | -0.32 | 0.18 | -0.32 | -0.12 | 0.00 | 0.12 | -0.24 | -0.45 | 0.34 |
| ILD1.5 | -0.31 | -0.31 | 0.50 | 0.40 | 0.02 | -0.01 | 0.04 | -0.43 | 0.35 | -0.29 | -0.03 |
| ILD1.9 | -0.10 | 0.36 | 0.32 | 0.09 | -0.54 | 0.01 | -0.15 | 0.16 | -0.06 | 0.08 | -0.63 |
| ILD2.5 | -0.08 | 0.65 | 0.12 | -0.13 | 0.55 | 0.27 | 0.04 | -0.07 | 0.07 | -0.36 | -0.13 |
| ILD2.8 | -0.42 | -0.08 | -0.56 | 0.09 | 0.23 | -0.14 | 0.15 | -0.10 | 0.26 | 0.25 | -0.50 |
| ILD3.7 | 0.41 | 0.14 | -0.32 | 0.38 | -0.05 | -0.09 | -0.54 | 0.06 | 0.46 | -0.22 | 0.02 |
| ILD3.9 | 0.12 | -0.29 | -0.14 | -0.25 | 0.05 | -0.11 | -0.34 | -0.39 | -0.47 | -0.42 | -0.37 |
| ILD4.6 | 0.44 | 0.22 | -0.17 | 0.34 | -0.18 | -0.11 | 0.62 | -0.37 | -0.16 | -0.09 | -0.07 |
| ILD4.8 | 0.14 | -0.33 | 0.00 | -0.11 | 0.00 | 0.03 | 0.37 | 0.59 | 0.20 | -0.52 | -0.25 |
| ILD5.7 | 0.00 | -0.13 | 0.08 | 0.65 | 0.39 | 0.12 | -0.10 | 0.33 | -0.49 | 0.11 | -0.10 |

**Table S6** | Eigen-analysis of **C**. Eigenvalues (λ) are bolded with 95% CI in italics; eigenvectors are beneath.

| | $e_1$ | $e_2$ | $e_3$ | $e_4$ | $e_5$ | $e_6$ | $e_7$ | $e_8$ | $e_9$ | $e_{10}$ | $e_{11}$ |
|---|---|---|---|---|---|---|---|---|---|---|---|
| λ | **14.31** | **6.75** | **4.23** | **2.39** | **0.26** | **-0.01** | **-0.11** | **-0.37** | **-0.93** | **-3.11** | **-17.74** |
| 95% CI | *0.51; 28.04* | *-4.10; 17.88* | *-0.37; 8.95* | *-7.76; 12.42* | *-3.45; 4.12* | *-0.28; 0.26* | *-0.63; 0.41* | *-5.01; 4.31* | *-3.22; 1.37* | *-11.02; 4.77* | *-36.21; 0.72* |
| Size | -0.06 | 0.28 | 0.81 | 0.15 | 0.16 | 0.02 | 0.06 | -0.23 | 0.34 | -0.13 | -0.14 |
| ILD1.2 | -0.49 | -0.19 | -0.03 | 0.07 | 0.17 | 0.04 | -0.61 | -0.04 | -0.02 | 0.21 | -0.52 |
| ILD1.5 | -0.27 | -0.32 | 0.16 | -0.62 | 0.41 | 0.05 | 0.06 | 0.22 | -0.02 | -0.39 | 0.18 |
| ILD1.9 | -0.25 | -0.45 | 0.17 | 0.14 | -0.44 | 0.47 | 0.28 | 0.29 | 0.26 | 0.18 | 0.01 |
| ILD2.5 | 0.71 | -0.16 | 0.15 | -0.09 | 0.11 | 0.18 | -0.13 | 0.39 | -0.05 | -0.02 | -0.47 |
| ILD2.8 | -0.18 | 0.33 | -0.13 | -0.31 | 0.12 | 0.14 | 0.59 | -0.05 | -0.16 | 0.28 | -0.50 |
| ILD3.7 | 0.11 | -0.19 | -0.26 | 0.28 | 0.67 | 0.02 | 0.19 | -0.02 | 0.48 | 0.30 | 0.12 |
| ILD3.9 | -0.03 | 0.14 | -0.14 | 0.33 | 0.22 | 0.72 | -0.02 | -0.17 | -0.31 | -0.40 | 0.05 |
| ILD4.6 | 0.04 | -0.14 | 0.39 | 0.05 | 0.20 | 0.05 | -0.01 | -0.02 | -0.60 | 0.56 | 0.33 |
| ILD4.8 | -0.04 | 0.58 | -0.04 | -0.26 | 0.00 | 0.32 | -0.34 | 0.40 | 0.26 | 0.28 | 0.28 |
| ILD5.7 | 0.25 | -0.19 | -0.04 | -0.46 | -0.14 | 0.31 | -0.15 | -0.69 | 0.21 | 0.18 | 0.05 |
